# AAV-based temporal APOE4-to-APOE2 replacement reveals rebound adaptation and RAB24-mediated Aβ and cholesterol dysregulation

**DOI:** 10.1101/2025.07.08.663694

**Authors:** Ruihan Yang, Yue Li, Xingchen Zhao, Lin Wang, Jialin Wu, Jian Sima

## Abstract

APOE-targeted gene therapy offers a promising strategy for modifying Alzheimer’s disease (AD) risk, yet the temporal dynamics and context-dependent effects of APOE isoform modulation remain poorly defined. Here, we developed a rapid AAV-based platform enabling inducible in vivo replacement of APOE4 with APOE2. In 5×FAD mice, sustained APOE4 expression exacerbated cognitive decline, Aβ deposition (parenchymal and vascular), and glial activation, whereas long-term APOE2 expression—with concurrent APOE4 silencing—significantly reversed these pathological features and rescued cognitive function. In contrast, short-term APOE2 replacement conferred no benefit and unexpectedly worsened behavioral and pathological outcomes. Transcriptomic profiling revealed that APOE4-associated gene signatures were broadly reversed by long-term APOE2 expression, but paradoxically aggravated by short-term replacement. Among these, RAB24—a regulator of autophagic trafficking—was upregulated by APOE4 and short-term APOE2 but suppressed by long-term APOE2. RAB24 elevation impaired Aβ clearance and cholesterol homeostasis via lysosomal retention in primary astrocytes and neurons. Together, these findings uncover a rebound-adaptation mechanism that shapes APOE2 therapeutic outcomes, identify RAB24 as a modifiable node in Aβ and cholesterol metabolism, and establish a temporally controlled gene therapy platform to inform the design of future APOE-targeted interventions in AD.

## Introduction

Alzheimer’s disease (AD), the most common form of dementia, is characterized by Aβ plaques, tau tangles, neuroinflammation, and cognitive decline, with the apolipoprotein E (APOE) gene standing out as the strongest genetic risk factor for late-onset AD^1,2^. Individuals carrying the *APOE* ε4 allele face a ∼3- to ∼14.5-fold increased risk of AD and an earlier age of onset^3,4^, while the rare ε2 allele confers protection and delays pathology^5,6^. These isoform-specific effects are attributed to differences in lipid metabolism, receptor binding, Aβ clearance, and immune signaling in the brain^7,8^.

Given the central role of APOE in AD pathogenesis, therapeutically targeting APOE—particularly replacing the pathogenic ε4 isoform with the neuroprotective ε2 variant—has become an area of active investigation. In preclinical studies, adeno-associated virus (AAV)-mediated delivery of APOE2 into the brain has shown the potential to reduce Aβ pathology and improve cognitive performance in mouse models^9–12^. Complementarily, targeted suppression of APOE4 using antisense oligonucleotides (ASOs) has been demonstrated to attenuate neurodegeneration and neuroinflammation in AD models^13,14^. These encouraging findings have catalyzed multiple translational efforts. Notably, Lexeo Therapeutics has initiated a Phase 1/2 clinical trial (LX1001) to evaluate intrathecal AAV-mediated APOE2 gene delivery in APOE4 homozygous individuals with early AD (NCT03634007). In parallel, other programs are developing miRNA-based platforms to selectively silence APOE4 expression^15^, while our recent study introduces an alternative AAV-based strategy involving silencing of mutant APP^swe^ mRNA alongside APOE2 overexpression as a combinatorial therapeutic approach^16^.

Despite this progress, several fundamental questions remain unresolved. It is unclear whether APOE2 supplementation alone is sufficient to counteract APOE4-driven pathology, or if co-suppression of APOE4 is essential for therapeutic benefit. Moreover, the timing of intervention—whether introduced early or late during disease progression— may critically determine efficacy. Additionally, the consequences of abrupt isoform switching in a disease-altered brain have yet to be systematically explored. A clearer understanding of how brain cells respond to dynamic APOE isoform transitions is vital to anticipate therapeutic outcomes and prevent maladaptive effects.

To address these knowledge gaps, we developed a rapid, inducible AAV-based platform that enables temporal control of APOE isoform switching in vivo, allowing APOE4-to-APOE2 replacement at defined stages in a disease context. Using the 5×FAD mouse model of Aβ pathology, we demonstrate that isoform switching reveals a rebound-adaptation mechanism that significantly impacts therapeutic outcomes. Furthermore, we identify RAB24, a cellular trafficking GTPase, as a key modulator of both Aβ clearance and cholesterol homeostasis. These findings underscore the importance of dosing schedule, cellular context, and replacement duration in optimizing APOE-targeted strategies for AD.

## Results

### Development of a dual AAV vector system for temporal control of APOE isoform expression

To systematically evaluate the timing and efficacy of APOE-targeted gene therapy, we designed a dual AAV vector system enabling inducible control of APOE isoform expression using a Cre-loxP recombination strategy (Fig. 1A). Because APOE is predominantly expressed in astrocytes, both AAV constructs employed the astrocyte-specific GfaABC1D promoter to mimic endogenous expression patterns. In the first vector, a Myc-tagged APOE4 sequence—terminated by a stop codon—was flanked by loxP sites and followed by an HA-tagged APOE2 sequence. The second vector encoded Cre-ERT2, a tamoxifen-inducible Cre recombinase, along with EGFP, separated by a T2A self-cleaving motif (Fig. 1A). In the absence of tamoxifen, astrocytes co-transduced with both AAV particles express APOE4. Upon tamoxifen administration, Cre-mediated recombination excises the APOE4 cassette, terminating its expression and enabling APOE2 expression instead.

**Fig. 1.**
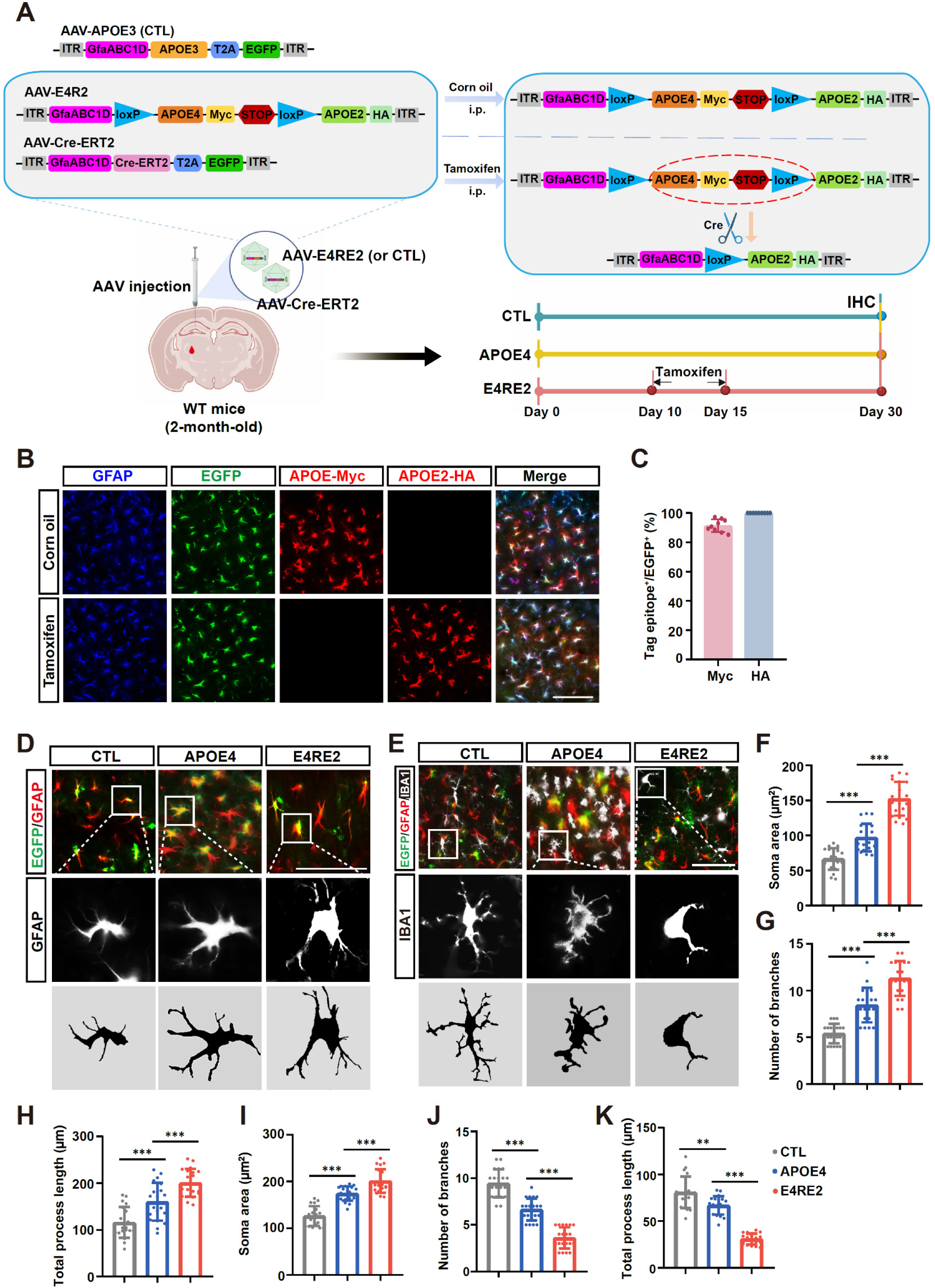
Inducible APOE4-to-APOE2 switching reveals glial activation following short-term replacement (**A**) Schematic of the dual AAV vector system and experimental timeline used to achieve tamoxifen-inducible APOE isoform switching in wild-type (WT) mice. AAV-mediated APOE3 expression served as the control (CTL). (**B**) Representative fluorescence images showing astrocyte-specific expression of EGFP, Myc-tagged APOE4, and HA-tagged APOE2 in the presence or absence of tamoxifen. (**C**) Quantification of EGFP⁺ cells co-expressing either APOE4-Myc or APOE2-HA. (**D, E**) Representative images and morphological tracings of GFAP⁺ astrocytes (D) and IBA1⁺ microglia (E) across CTL, APOE4, and E4RE2 treatment groups. (**F–H**) Quantification of astrocyte hypertrophy including soma area (F), number of branches (G), and total process length (H) in EGFP⁺/GFAP⁺ cells. (**I–K**) Quantification of microglial morphology, including soma area (I), branch number (J), and total process length (K). Data are presented as mean ± SD. Statistical significance determined by one-way ANOVA followed by Tukey’s post hoc test. \**P* < 0.05, \*\**P* < 0.01, \*\*\**P* < 0.001. Scale bar, 50 µm.

To validate this system, we packaged the two constructs into AAV5 serotype particles, known for their high efficiency in astrocyte transduction^17^. Following intrahippocampal injection, robust EGFP expression confirmed widespread viral transduction throughout the hippocampus (Fig. S1). Co-immunolabeling with GFAP and APOE4-Myc demonstrated strong astrocyte specificity and effective dual-vector delivery (Fig. 1B, C). Tag-specific immunostaining revealed isoform-selective expression: APOE4 (Myc-tagged) was prominently expressed in the absence of tamoxifen, whereas tamoxifen administration triggered Cre-mediated recombination, leading to APOE4 silencing and induction of APOE2 (HA-tagged) expression (Fig. 1B, C). These results confirm the functionality of the system for temporally controlled and astrocyte-specific modulation of APOE isoform expression in vivo.

### Two-week APOE4-to-APOE2 replacement induces unexpected glial activation in wild-type mice

To investigate the cellular consequences of APOE isoform replacement in the absence of Aβ pathology, we employed our inducible AAV system in wild-type mice. Two-month-old (Day 0) mice received intrahippocampal injections of AAV5 particles encoding APOE4 and Cre-ERT2, followed by daily tamoxifen administration for 5 consecutive days beginning 10 days post-injection to initiate APOE4-to-APOE2 replacement (hereafter referred to as E4RE2, APOE4 replaced by APOE2). On day 30, brains were collected for immunohistochemical (IHC) analysis (Fig. 1A). Compared to APOE3-expressing controls, APOE4 expression in astrocytes induced significant GFAP^+^ astrocyte hypertrophy, characterized by increased soma area, branching complexity, and process length (Fig. 1D, F-H). In parallel, nearby Iba1^+^ microglia exhibited increased density and morphological features consistent with reactive activation (Fig. 1E, I-K). Surprisingly, APOE2 replacement further amplified both astrocytic and microglial activation beyond levels observed with sustained APOE4 expression (Fig. 1D-K). These changes occurred in the absence of Aβ deposition or neuronal loss, indicating that acute isoform switching is sufficient to perturb glial homeostasis.

Together, these findings reveal a previously unrecognized rebound-like response to APOE isoform transition, highlighting the importance of dosing strategy and timing in the design of APOE-targeted therapies.

### Long-term, but not short-term, E4RE2 confers cognitive protection in 5×FAD mice

To evaluate the therapeutic impact of APOE isoform replacement on Alzheimer’s pathology, we utilized our inducible AAV system in 5×FAD mice and assessed behavioral outcomes following short-term versus long-term APOE4-to-APOE2 replacement (E4RE2). Two-month-old mice received bilateral injections of AAV5 viral particles encoding APOE4 or APOE3 (CTL), along with Cre-ERT2 into both the hippocampus and cortex. Tamoxifen-induced switching to APOE2 was initiated either at 3 months of age (long-term E4RE2 group) or at 5.3 months of age (short-term E4RE2 group), with all animals aged to 6 months before behavioral testing and following immunohistochemical (IHC) analyses (Fig. 2A).

**Fig. 2.**
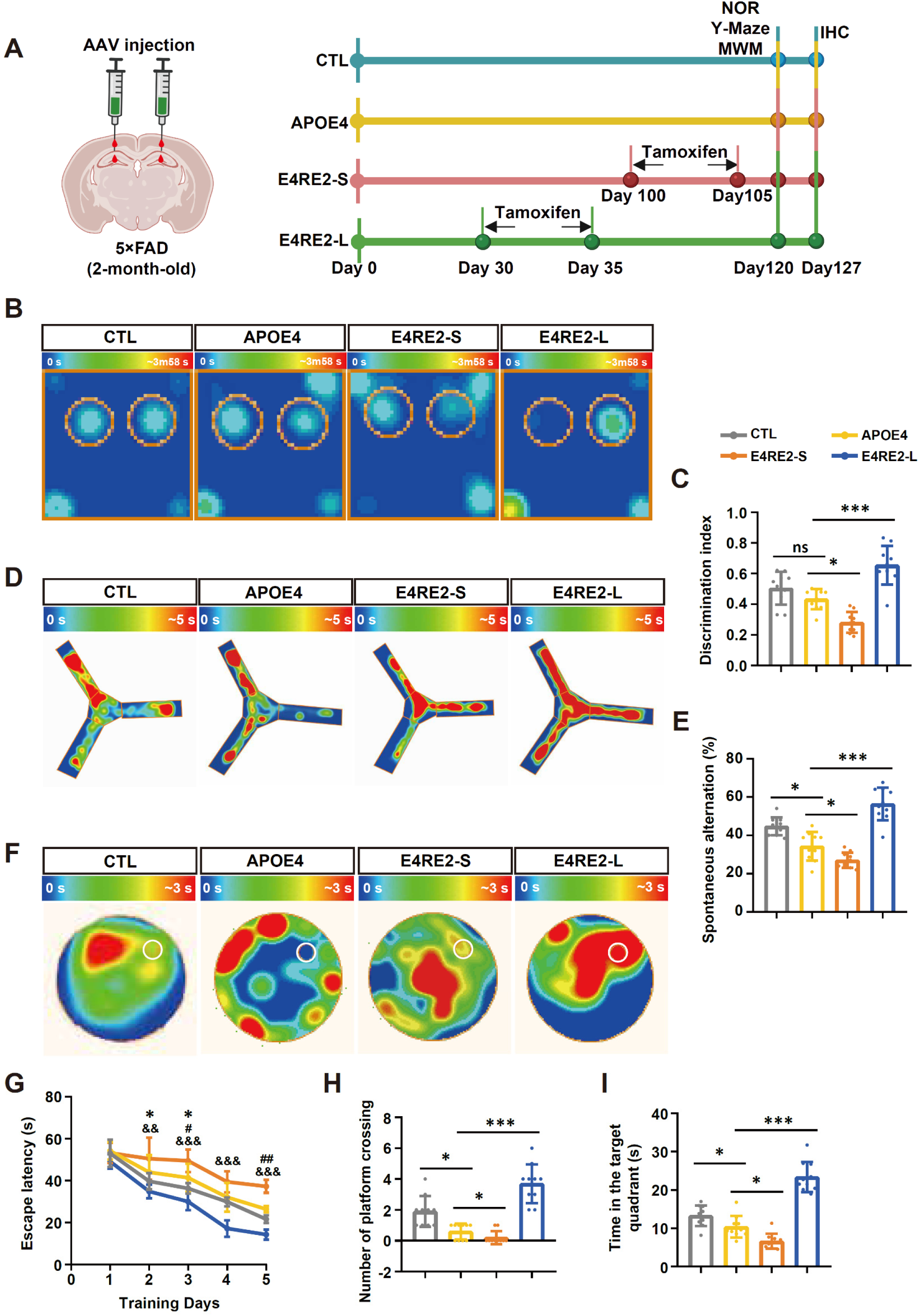
Long-term, but not short-term, E4RE2 improves cognitive function in 5×FAD mice (**A**) Schematic of the experimental design. Two-month-old 5×FAD mice received bilateral AAV injections into the hippocampus and cortex. Tamoxifen was administered either at 3 months of age (long-term replacement, E4RE2-L) or at 5.3 months (short-term replacement, E4RE2-S) to induce APOE4-to-APOE2 switching. Behavioral testing was conducted at 6 months of age. (**B, C**) Novel Object Recognition (NOR) test: representative heatmaps (B) and discrimination index (C) across all four groups. (**D, E**) Y-maze test: representative exploration heatmaps (D) and quantification of spontaneous alternation percentage (E). (**F-I**) Morris Water Maze (MWM): (F) representative heatmaps showing visit frequency in the target quadrant during the probe trial (Day 6); (G) escape latency during 5 days of training; (H) time spent in the target quadrant; (I) number of platform crossings during the probe trial. Statistical symbols in (G): * indicates APOE4 vs. CTL; # indicates E4RE2-S vs. APOE4; & indicates E4RE2-L vs. APOE4. Data are presented as mean ± SD. Statistical comparisons were performed using one-way ANOVA (C, E, H, I) or two-way ANOVA (G), followed by Tukey’s HSD post hoc test. *p < 0.05, **p < 0.01, ***p < 0.001; ns, not significant.

Cognitive performance was assessed using the novel object recognition (NOR) task, Y-maze, and Morris water maze (MWM). As expected, APOE4-expressing mice exhibited significant impairments in spatial memory and learning compared to APOE3-expressing controls in the Y-maze and MWM tests (Fig. 2D-I). While NOR performance did not reach statistical significance, the APOE4 group showed a trend toward a reduced discrimination index (Fig. 2B, C). Notably, long-term E4RE2 significantly rescued cognitive deficits across all three behavioral paradigms, indicating robust neuroprotection. In contrast, short-term E4RE2 conferred no benefit and, unexpectedly, led to greater impairments than sustained APOE4 expression (Fig. 2B-I), suggesting that abrupt isoform switching may exacerbate functional decline. These findings highlight the importance of timing and duration in APOE-targeted interventions for AD.

### Aβ burden in brain parenchyma and vasculature correlates with cognitive performance across E4RE2 regimens

To investigate whether the divergent cognitive outcomes observed in long-versus short-term E4RE2-treated mice were associated with differences in Aβ pathology, we conducted IHC analyses of amyloid deposition in both the brain parenchyma and vasculature. Aβ immunolabeling revealed pronounced plaque accumulation in the hippocampus and cortex of APOE4-expressing 5×FAD mice, as expected (Fig. 3A, B). Notably, these mice—but not age-matched 5×FAD controls expressing exogenous APOE3—also exhibited substantial Aβ deposition within the blood vessels of both the brain parenchyma and meninges (Fig. 3C, D), consistent with previous findings implicating APOE4 in promoting cerebral amyloid angiopathy^18^. Long-term E4RE2 treatment significantly reduced both parenchymal and meningeal Aβ burden relative to sustained APOE4 expression (Fig. 3A-H). By contrast, short-term E4RE2 led to further exacerbation of Aβ deposition in both compartments, with particularly prominent accumulation along cerebrovascular and meningeal surfaces (Fig. 3C, D, G and H). These findings suggest that abrupt APOE isoform replacement may transiently impair peripheral Aβ clearance or disrupt vascular trafficking. Importantly, the extent and distribution of Aβ pathology across treatment groups closely mirrored their behavioral phenotypes.

**Fig. 3.**
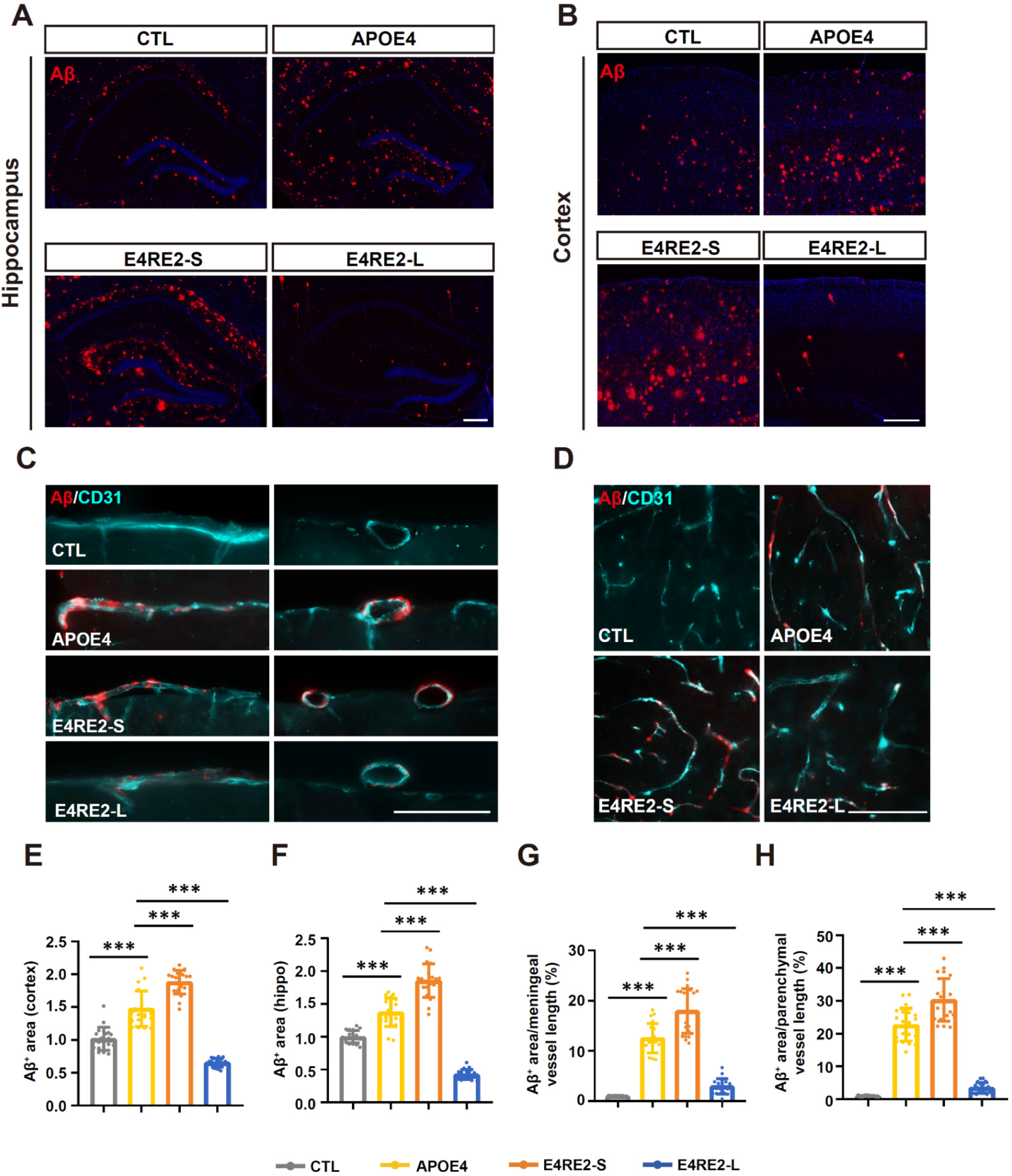
E4RE2-S exacerbates Aβ deposition in both brain parenchyma and vasculature compared to sustained APOE4 expression (**A, B)** Representative IHC images showing Aβ plaques (red) in the hippocampus (A) and cortex (B) across treatment groups. (**C**) Representative IHC images showing co-labeling of Aβ (red) and CD31 (blue), an endothelial cell and vascular marker, in the meninges from both coronal (left) and sagittal (right) brain sections. (**D**) Co-labeling of Aβ and CD31 in cortical parenchyma. (**E, F**) Quantification of Aβ plaque burden in the hippocampus (E) and cortex (F), expressed as plaque area relative to the CTL group (normalized to 1.0). (**G, H**) Quantification of vascular Aβ deposition in the meninges (G) and cortex (H), calculated as Aβ-positive area per 10 μm of CD31⁺ vascular length (%). Data are presented as mean ± SD. Statistical comparisons were performed using one-way ANOVA followed by Tukey’s HSD post hoc test. ***p < 0.001. Scale bar, 100 μm.

### Glial remodeling reveals rebound-like activation following short-term APOE isoform replacement

To explore whether glial cells contribute to the divergent outcomes observed in long-versus short-term E4RE2, we performed detailed morphological analyses of astrocytes and microglia surrounding Aβ plaques across treatment groups. In APOE4-expressing 5×FAD mice, compared to APOE3-expressing controls, GFAP immunostaining combined with Sholl analysis revealed marked astrocytic hypertrophy in both hippocampal and cortical regions, characterized by enlarged somata, increased process branching, and extended arborization (Fig. 4A, B). Microglia labeled by IBA1 also exhibited a reactive phenotype, with elevated cell density, amoeboid morphology, and increased CD68 immunoreactivity (Fig. 4C-E, Fig. S2), consistent with pro-inflammatory activation and APOE4-driven microglial chemotaxis^19^. Long-term E4RE2 markedly attenuated these glial responses, aligning with reduced Aβ pathology and improved cognitive performance. In contrast, short-term E4RE2 not only failed to reverse glial response but further exacerbated astrocytic and microglial activation (Fig. 4A–E, Fig. S2). We further assessed microglial morphology in perivascular regions. Similar to plaque-associated microglia, cells adjacent to Aβ-laden vasculature in APOE4 and short-term E4RE2 groups exhibited amoeboid morphology with retracted, simplified processes—phenotypes that were normalized by long-term E4RE2 treatment (Fig. 4F–I).

**Fig. 4.**
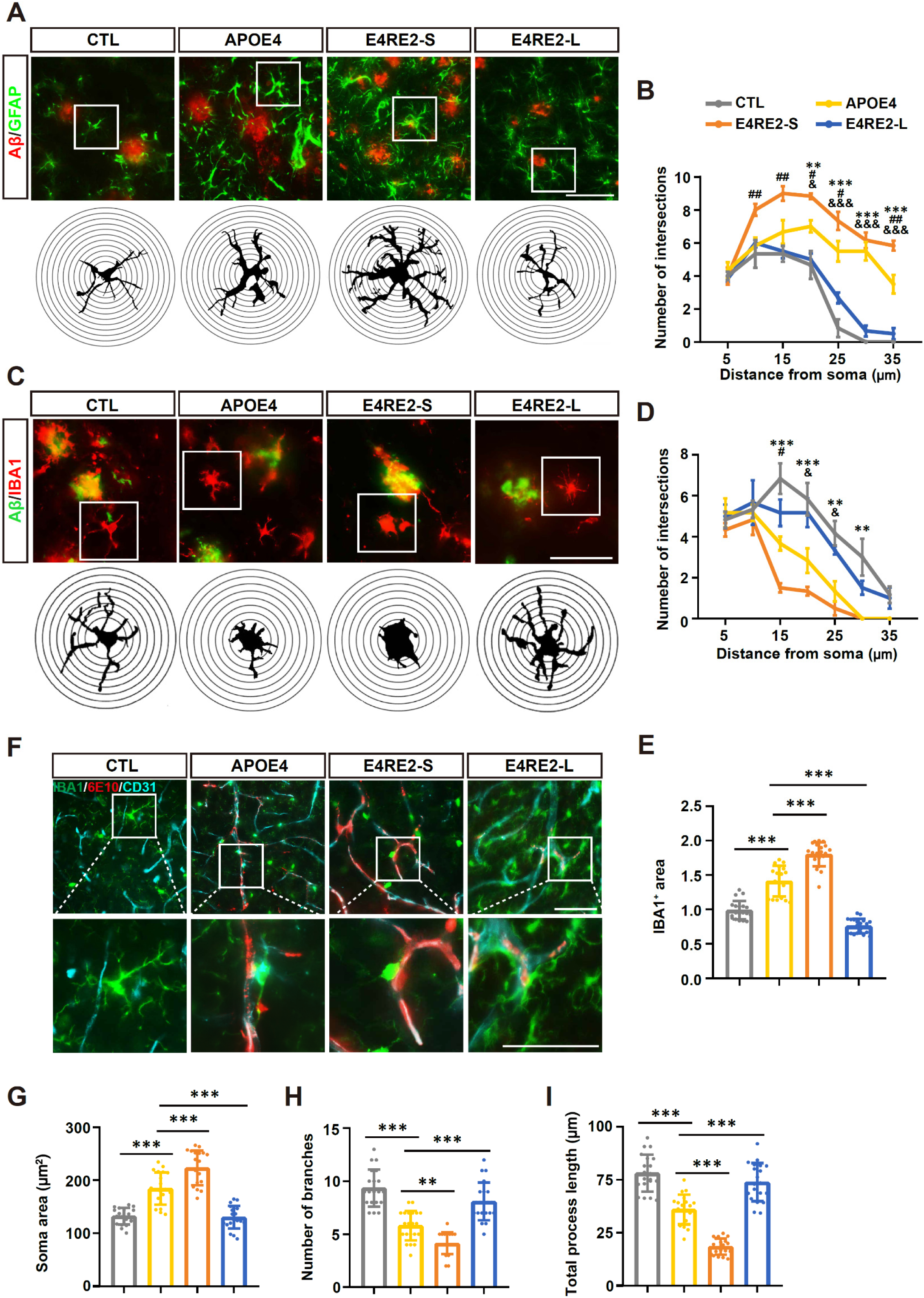
Glial morphological analyses reveal exacerbated astrocytic and microglial activation following E4RE2-S (**A**) Representative IHC images of GFAP^+^ astrocytes adjacent to Aβ plaques (top) and corresponding traced reconstructions of astrocyte morphology (bottom), across the four experimental groups. (**B**) Sholl analysis quantifying astrocytic process complexity. (**C, D**) As in (A, B), but for IBA1⁺ microglia: representative morphologies (C) and Sholl-based complexity analysis (D). (**E**) Quantification of IBA1^+^ area in the hippocampus. (**F**) Representative IHC images showing triple labeling of Aβ (6E10, red), microglia (Iba1, green), and vasculature (CD31, blue) in the cortex. (**G-I**) Quantification of microglial soma area (G), number of process branches (H), and total process length (I). Statistical symbols in Sholl analysis: * indicates APOE4 vs. CTL; # indicates E4RE2-S vs. APOE4; & indicates E4RE2-L vs. APOE4. Data are presented as mean ± SD. Statistical significance was determined using one-way ANOVA (E, G-I) or two-way ANOVA (B, D) followed by Tukey’s HSD post hoc test. **p < 0.01, ***p < 0.001. Scale bar, 50 μm.

These findings suggest that abrupt isoform switching provokes a rebound-like glial response. This heightened glial activation, which paralleled enhanced vascular Aβ deposition, implicates disrupted glial–vascular homeostasis in the pathological cascade and underscores the risk of unintended pro-inflammatory consequences following abrupt APOE modulation.

### Transcriptomic profiling reveals divergent molecular responses to short- and long-term E4RE2

To elucidate the molecular mechanisms underlying the distinct outcomes of short-versus long-term E4RE2, we performed bulk RNA sequencing on hippocampal tissues from 5×FAD mice expressing APOE4, short-term or long-term E4RE2, and APOE3-expressing controls. Compared to controls, APOE4-expressing 5×FAD mice exhibited widespread transcriptional dysregulation, with approximately 380 differentially expressed genes (DEGs). Long-term E4RE2 corrected a substantial subset of these APOE4-associated changes, with ∼1,000 DEGs relative to APOE4 mice. In contrast, short-term E4RE2 induced a broader and more pronounced shift in gene expression compared to APOE4, indicative of a widespread stress- or adaptation-like response following abrupt APOE isoform switching (Fig. 5A).

**Fig. 5.**
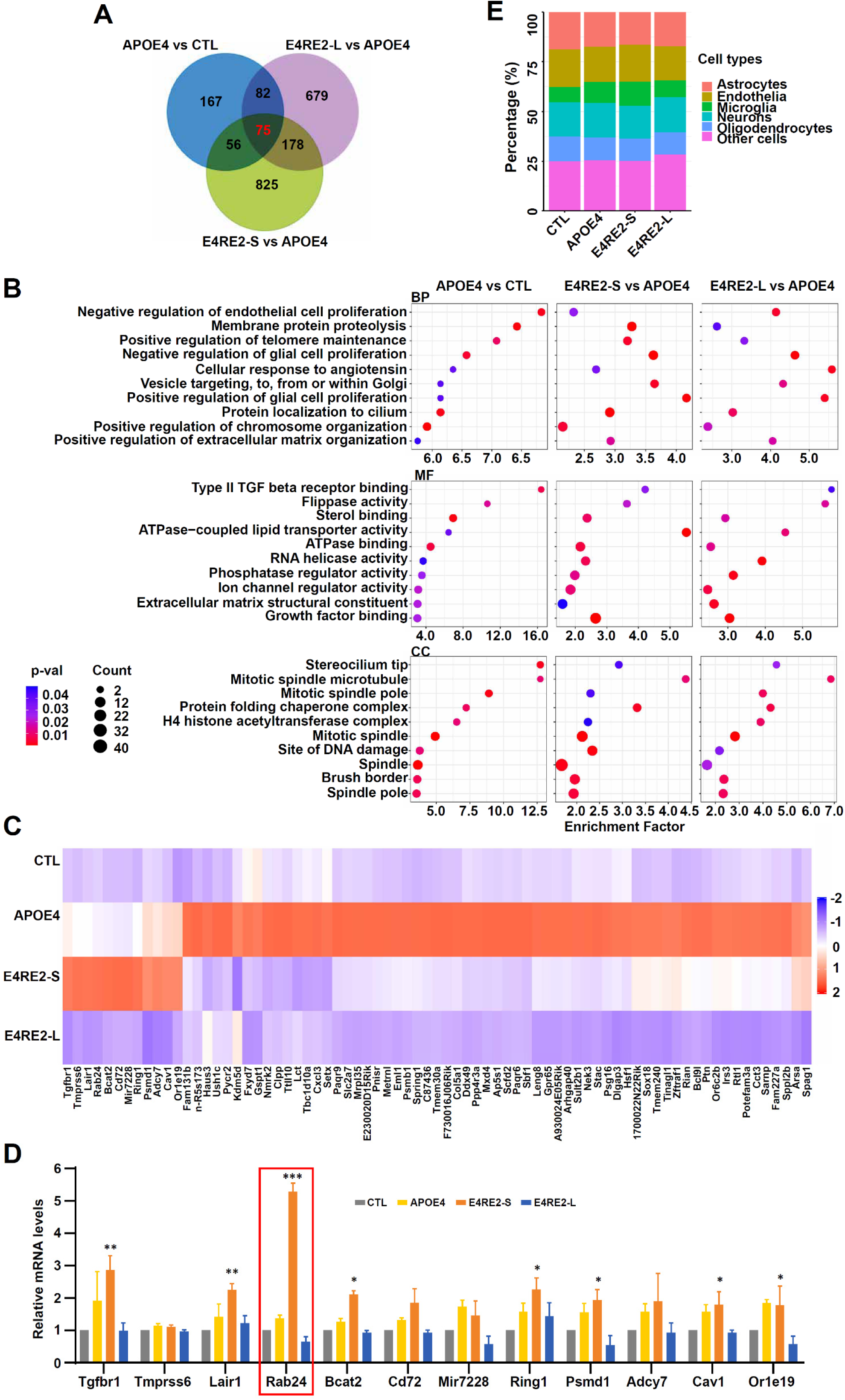
Transcriptomic profiling reveals distinct molecular responses to E4RE2-S and E4RE2-L (**A**) Venn diagram illustrating differentially expressed genes (DEGs) shared and uniquely regulated by APOE4, E4RE2-S and E4RE2-L. (**B**) Bubble plots of significantly enriched Gene Ontology (GO) terms—including Biological Process (BP), Molecular Function (MF), and Cellular Component (CC)—among DEGs from the indicated comparisons. (**C**) Heatmap displaying a set of 75 representative genes concordantly regulated across treatment groups, highlighting shared transcriptional signatures. (**D**) qPCR validation of 12 genes exhibiting rebound-like upregulation specifically in the E4RE2-S group. (**E**) Cell type deconvolution of bulk RNA-seq data showing estimated proportions of major brain cell types across experimental groups. Data in (D) are presented as mean ± SD. Statistical analysis was performed using one-way ANOVA followed by Tukey’s HSD post hoc test. * indicates E4RE2-S vs. APOE4; *p < 0.05, **p < 0.01, ***p < 0.001.

Gene Ontology (GO) enrichment analysis revealed significant overrepresentation of DEGs in pathways related to endothelial and glial proliferation, extracellular matrix remodeling, growth factor binding, mitotic spindle dynamics, and angiotensin signaling, spanning biological process (BP), molecular function (MF), and cellular component (CC) categories (Fig. 5B). A core set of 75 genes concordantly regulated across treatment groups was visualized by heatmap (Fig. 5C). Among these, 12 genes were uniquely exacerbated in the short-term E4RE2 group compared to APOE4, but reversed by long-term replacement (Fig. 5C). qPCR validation identified Rab24 as the most strongly upregulated gene in this subset (Fig. 5D). To explore potential shifts in brain cell-type composition, we performed transcriptomic deconvolution of the bulk RNA-seq data. This analysis revealed a significant increase in the inferred proportion of microglia following short-term E4RE2 treatment (Fig. 5E), consistent with the rebound-like glial activation observed histologically.

### RAB24 impairs Aβ clearance and cholesterol trafficking by promoting lysosomal retention

To investigate the cellular mechanisms underlying impaired Aβ clearance in APOE4-expressing and short-term E4RE2-treated mice, we focused on RAB24, a small GTPase involved in intracellular trafficking, identified as the most strongly upregulated gene by short-term E4RE2 treatment (Fig. 5D). To assess cell-type specificity, we measured RAB24 expression in primary neurons, astrocytes, and microglia. qPCR revealed robust expression in neurons and astrocytes, but minimal levels in microglia (Fig. S3), consistent with publicly available transcriptomic datasets (The human protein atlas).

To determine whether RAB24 modulates Aβ handling, we overexpressed RAB24 in primary astrocyte and neuron cultures exposed to oligomeric Aβ and performed cellular localization and clearance assays as previously established^20^. RAB24-overexpressing astrocytes and neurons exhibited pronounced intracellular accumulation of Aβ within LAMP1⁺ lysosomal compartments, as shown by immunofluorescence (Fig. 6A, B, G, H).

**Fig. 6.**
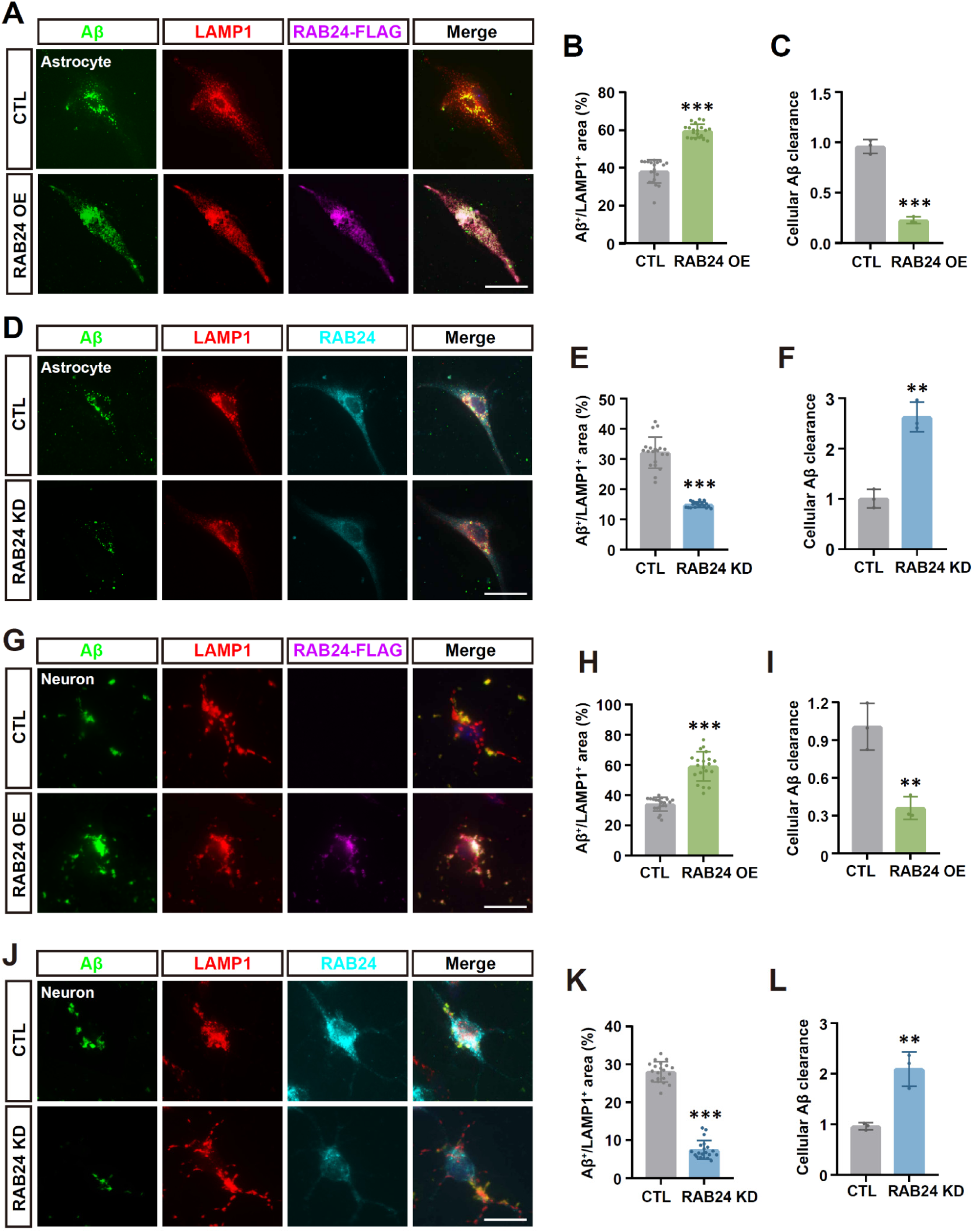
RAB24 disrupts Aβ degradation by promoting lysosomal sequestration (**A**) Representative images of primary astrocytes showing FAM-labeled Aβ oligomers (green), LAMP1⁺ lysosomes (red), and Flag-tagged RAB24 (purple) under empty vector control (top, CTL) and RAB24 overexpression (bottom, OE) conditions. (**B**) Quantification of Aβ⁺ area co-localized with LAMP1⁺ puncta in astrocytes. (**C**) ELISA quantification of Aβ clearance in astrocytes with or without RAB24 overexpression. (**D-F**) Same analyses as in (A–C), comparing control scrambled siRNA control (CTL) and RAB24 knockdown (KD) via siRNA. (**G–I**) As in (A–C), but performed in primary cortical neurons. (**J–L**) As in (D– F), but performed in neurons. Data are as mean ± SD. Statistical analysis was performed using student’s t-test. *p < 0.05, **p < 0.01, ***p < 0.001. Scale bar, 20 µm.

ELISA quantification confirmed impaired Aβ clearance relative to control-transduced cells (Fig. 6C, I). Conversely, shRNA-mediated knockdown of RAB24 significantly enhanced Aβ degradation and reduced lysosomal retention in both neurons and astrocytes (Fig. 6D-F, J-L).

Given recent evidence linking APOE4 to cholesterol dysregulation in astrocytes^21^, we further investigated RAB24 function in cholesterol regulation. Filipin staining revealed a marked increase (∼25.02%) in lysosomal cholesterol accumulation in RAB24-overexpressing astrocytes compared to controls (Fig. 7A, B), while RAB24 knockdown significantly reduced cholesterol intensity within LAMP1⁺ compartments (Fig. 7C, D). These results implicate RAB24 as a critical regulator of both Aβ and cholesterol trafficking, functioning by sequestering cargo within lysosomes and thereby impeding their clearance or recycling.

**Fig. 7.**
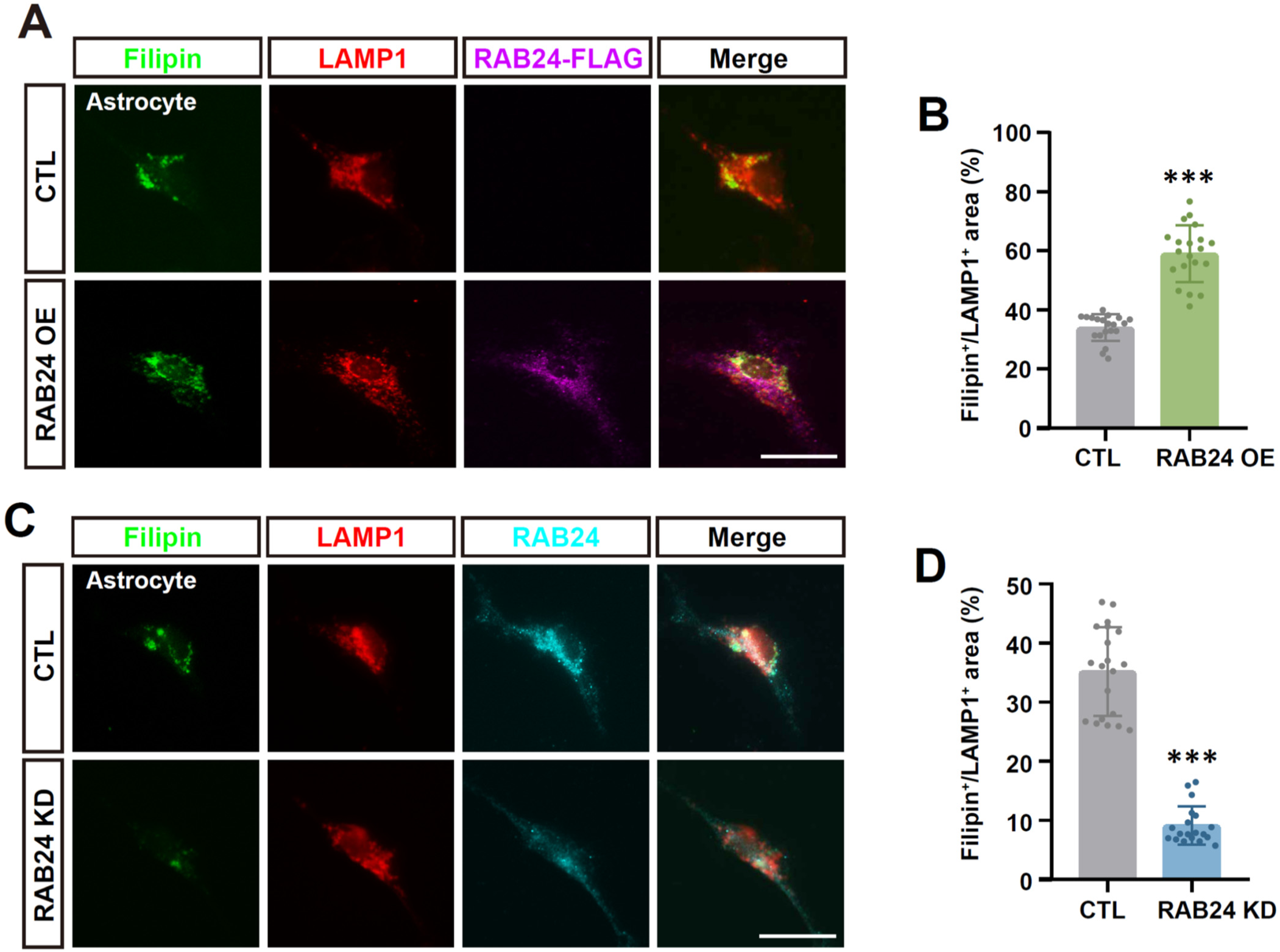
RAB24 impairs cholesterol homeostasis via lysosomal trapping (**A**) Representative images of primary astrocytes stained for cholesterol (Filipin, green), lysosomes (LAMP1, red), and Flag-tagged RAB24 (purple), under empty vector control (top, CTL) and RAB24 overexpression (bottom, OE) conditions. (**B**) Quantification of Filipin⁺ area co-localized with LAMP1⁺ lysosomes in astrocytes. (**C, D**) Same analyses as in (A, B), but comparing scrambled siRNA control (CTL) and RAB24 knockdown (KD) via siRNA. Data are presented as mean ± SD. Statistical significance was determined using student’s t-test. *p < 0.05, **p < 0.01, ***p < 0.001. Scale bar, 20 μm.

Together, these findings identify RAB24 as a dual-pathway negative regulator of Aβ degradation and cholesterol homeostasis. Its downregulation by long-term APOE2 replacement suggests that modulation of RAB24-mediated trafficking may contribute to the therapeutic benefits of APOE-targeted intervention in AD.

## Discussion

The apolipoprotein E (APOE) genotype remains the most potent genetic determinant of Alzheimer’s disease (AD) risk, with APOE4 conferring up to a 14.5-fold increase in late-onset AD incidence, while APOE2 is associated with significantly reduced risk and delayed onset^2,4,8,22^. These opposing isoform effects have sparked intense interest in therapeutic APOE modulation^23,24^. However, despite promising advances in gene-based approaches, key questions remain about the timing, duration, and physiological consequences of isoform switching within the diseased brain—issues that are particularly relevant to clinical translation. Here, we developed a rapid, AAV-based platform enabling temporally controlled replacement of APOE4 with APOE2 in vivo. Our results demonstrate that the therapeutic impact of isoform switching is critically shaped not only by the presence of APOE2, but also by the duration of expression and the underlying pathophysiological context.

Consistent with recent studies linking APOE4 to enhanced amyloidogenic processing, impaired clearance, and vascular dysfunction^24–26^, sustained APOE4 expression in the 5×FAD model exacerbated hallmark AD phenotypes, including amyloid-β (Aβ) accumulation, glial activation, and cognitive decline. In striking contrast, long-term APOE2 expression—introduced with concurrent APOE4 silencing—reversed these pathological features, significantly improving both behavioral performance and reducing parenchymal and vascular Aβ burden (Fig. 2–4). These observations align with earlier reports that APOE2 promotes Aβ clearance, supports lipid transport, and enhances neuronal resilience in both murine and iPSC-derived models^12,27–29^. Moreover, human carriers of APOE2/3 or APOE2/2 genotypes show preserved cognitive function and lower plaque load even in the presence of aging or familial AD mutations^30–32^.

Unexpectedly, short-term APOE2 replacement not only failed to confer benefit but worsened pathology, inducing increased vascular Aβ deposition and heightened glial reactivity (Fig. 2–4). Notably, this rebound-like effect occurred despite effective APOE4 silencing, suggesting that abrupt isoform switching may disrupt glial–vascular homeostasis and trigger maladaptive responses. These findings have important clinical implications. APOE isoforms are known to influence Aβ immunotherapy outcomes, including susceptibility to amyloid-related imaging abnormalities (ARIA)^33,34^. Our results raise the possibility that APOE-based gene therapies, if administered abruptly or at inappropriate stages of disease, may amplify ARIA risk or provoke neuroinflammatory cascades, particularly in aged or vulnerable individuals^35,36^.

To dissect the underlying molecular responses, we conducted transcriptomic profiling of brain tissue. Long-term APOE2 expression broadly reversed APOE4-induced signatures, including those linked to gliosis, extracellular matrix remodeling, and vesicular trafficking. However, short-term APOE2 replacement aggravated a distinct subset of these pathways, pointing to a rebound-adaptation program that may override the intended therapeutic effect. Among the top hits, we identified RAB24, a GTPase involved in vesicular transport and autophagy machinery^37^, as markedly elevated under APOE4 expression and further upregulated by short-term APOE2 treatment (Fig. 5C, D). Mechanistically, RAB24 overexpression in primary astrocytes impaired Aβ clearance and cholesterol trafficking by trapping them within LAMP1⁺ lysosomal compartments, a phenotype rescued by shRNA-mediated knockdown (Fig. 6-7). These results position RAB24 as a bottleneck in endolysosomal trafficking that could be co-targeted to enhance the efficacy of APOE2-based interventions.

Notably, our findings echo emerging transcriptomic and lipidomic data from a recent abstract presentation titled *“Mid-life APOE4 to APOE2 ‘Switching’ Alters the Cerebral Transcriptome and Lipidome in a Transgenic Mouse Model”* (AlzForum, 2023). That study, based on a germline model, reported widespread remodeling in response to isoform switching. In contrast, our platform offers rapid and precise temporal control, closely mimicking AAV gene therapy applications. Although full data from that transgenic model remain unpublished, the convergence in transcriptomic remodeling supports our conclusion that timing and isoform context are key determinants of APOE therapeutic outcomes.

In summary, our study reveals that the timing and dynamics of APOE isoform replacement are crucial in determining therapeutic efficacy, with long-term APOE2 expression conferring benefit, and short-term switching inducing rebound pathology. We identify RAB24 as a mechanistic link between APOE context and impaired Aβ/cholesterol trafficking via lysosomal trapping. While informative, our study has several limitations: we employed an AAV serotype distinct from clinically preferred, BBB-penetrant variants, and the 5×FAD model exhibits aggressive Aβ pathology that may not fully reflect sporadic AD. Further studies are warranted to validate RAB24 function within the cerebrovascular system and to investigate its broader involvement in peripheral Aβ clearance and lipid/cholesterol metabolism—pathways central to APOE-related AD pathogenesis^21^. Nonetheless, these insights deepen our understanding of APOE biology in AD and offer a mechanistic framework for rationally designing APOE-targeted interventions— emphasizing not only the ‘what’ (isoform) but the ‘when’ and ‘how’ of gene therapy delivery.

## Materials and methods

### AAV preparation

To generate the AAV-E4RE2 vector, the pAAV-CAG-EGFP plasmid (Addgene, #37825) was modified by inserting a T2A self-cleaving peptide and a multiple cloning site (MCS) downstream of the EGFP sequence, resulting in the pAAV-CAG-EGFP-T2A-MCS backbone. The ubiquitous CAG promoter was subsequently replaced with the astrocyte-specific GfaABC1D promoter. Human APOE4 cDNA, fused to a Myc tag and containing a stop codon, was flanked by loxP sites at both the 5′ and 3′ ends via PCR cloning. Downstream of the 3′ loxP site, APOE2 cDNA tagged with an HA epitope at the C-terminus was inserted. For the AAV-CreERT2 and AAV-APOE3 vectors, the CAG-EGFP cassette was replaced with the GfaABC1D promoter driving CreERT2 or APOE3 expression. An EGFP reporter was inserted into the MCS downstream of a T2A motif to enable bicistronic expression.

Recombinant AAV vectors were produced and purified according to protocols provided by Addgene, with modifications optimized in-house. Briefly, HEK293T cells were co-transfected with the AAV expression plasmid along with the necessary helper plasmids to support AAV5 replication and packaging. After 72 h, transfected cells were harvested and subjected to three freeze–thaw cycles to lyse the cells. The resulting lysates were clarified by centrifugation, and viral particles were purified by iodixanol step-gradient ultracentrifugation. Purified viral preparations were concentrated, and genomic titers were determined by quantitative PCR using primers targeting the viral ITR sequence. Aliquots were stored at –80 °C until use.

### Mice

In-house C57BL/6J and transgenic 5×FAD mice (JAX, strain no. 034848) were used. For AAV experiments, two-month-old mice (male:female ratio ≈ 1:1) were randomly assigned to treatment groups. All animals were housed under standard conditions (25 ± 2 °C, 55% relative humidity, 12 h light/dark cycle) with free access to food and water. For APOE isoform switching, tamoxifen (Sigma) was dissolved in corn oil and administered via intraperitoneal injection at 100 mg/kg once daily for 5 consecutive days, starting at designated time points post-AAV injection as indicated in the experimental timeline. Control animals received equal volumes of corn oil without tamoxifen. All experimental procedures were approved by the Laboratory Animal Care Committee of China Pharmaceutical University and conducted in accordance with national guidelines for the care and use of laboratory animals.

### Brain stereotaxic injection

Mice were anesthetized via intraperitoneal injection of 1% pentobarbital sodium (50 mg/kg) and secured in a stereotaxic apparatus. For AAV administration, 2 μL of viral suspension was injected unilaterally or bilaterally into either the hippocampal CA1 region (anteroposterior −2.2 mm, mediolateral ±1.7 mm, dorsoventral −2.4 mm from bregma) or cortical layers II–IV (dorsoventral −0.4 mm) using a microsyringe pump at a rate of 0.25 μL/min. Following injection, the needle was left in place for 10 min to prevent backflow and then withdrawn slowly. For APOE isoform replacement experiments, AAV-APOE3 (control) or AAV-E4RE2 (2 × 10¹² vg/mL) were premixed with AAV-CreERT2 (2 × 10¹² vg/mL) at a 1:1 volume ratio and injected as described above. Mice were placed on a thermostatically controlled heating pad and closely monitored until full recovery.

### Behavioral testing

Cognitive function was evaluated using the novel object recognition (NOR), Morris water maze (MWM), and Y-maze spontaneous alternation tasks. For the NOR test, mice were habituated to an open field arena (45 × 45 × 40 cm) for 2 min. After a 30-min inter-trial interval, they explored two identical objects for 8 min (training). Following a 1 h delay, one object was replaced with a novel item, and mice explored for another 8 min (test). Exploration time was recorded and the discrimination index (DI) calculated as: DI = T_Novel / (T_Novel + T_Familiar).

The MWM was used to assess spatial learning and memory. Mice were trained to locate a submerged escape platform (10 cm diameter, 1 cm below the surface) in a circular pool (130 cm diameter, 60 cm deep) filled with opaque water maintained at 25 ± 1 °C. Training consisted of four trials per day over five consecutive days. Mice were released from different quadrants in a pseudorandom order and allowed 60 s to locate the platform. If unsuccessful, they were gently guided to the platform and allowed to remain there for 15 s to encode spatial cues. Inter-trial intervals were 15 s. On day 6, a 60 s probe trial was conducted with the platform removed, and the time spent in each quadrant was recorded. Swim paths were analysed using an automated tracking system.

The Y-maze spontaneous alternation test was used to assess short-term spatial working memory. The maze consisted of three arms (40 cm long × 6.8 cm wide × 15.5 cm high) arranged at 120° angles. Mice were placed at the end of one arm and allowed to explore freely for 8 min. An arm entry was counted when all four limbs entered an arm. Spontaneous alternation was defined as successive entries into all three arms (e.g., ABC, BCA). The percentage of alternation was calculated as: % Alternation = [(Number of alternations) / (Total arm entries − 2)] × 100. The maze was cleaned with 70% ethanol between trials to eliminate olfactory cues.

### Immunofluorescence

For tissue immunofluorescence, mice were euthanized and perfused with PBS followed by 4% paraformaldehyde (PFA). Brains were post-fixed in 4% PFA overnight, cryoprotected in 30% sucrose, embedded in OCT, and sectioned coronally at 20 μm using a cryostat (Leica CM1950). Sections were permeabilized in 0.3% Triton X-100 (3 × 10 min), blocked in 2.5% bovine serum albumin (BSA) for 1 h at room temperature (RT), and incubated with primary antibodies overnight at 4 °C. After PBS washes, sections were incubated with fluorophore-conjugated secondary antibodies for 1 h at RT, washed again, and mounted using DAPI-containing antifade medium. Images were acquired using a fluorescence microscope and analyzed with ImageJ. For cultured cells, coverslips were fixed in 4% PFA for 20 min at RT, permeabilized with 0.2% Triton X-100 for 20 min, and blocked with 2.5% BSA in PBS for 1 h. Cells were incubated with primary antibodies overnight at 4 °C, followed by incubation with appropriate secondary antibodies for 1 h at RT. Coverslips were mounted with antifade medium containing DAPI. Antibodies are listed in Supplementary Table 1.

### Analysis of glial morphology and Aβ pathology

Fluorescence images were captured using an Olympus IX73 fluorescence microscope. Morphological features, including soma area, total process length, and branch number, were quantified using ImageJ software with the Sholl Analysis plugin. For in vivo analysis of glial morphology, approximately 20 image fields—with or without Aβ plaque presence (from WT or 5×FAD mice)—were selected from cortical cryosections (20 μm thick) obtained from at least four mice per group. EGFP-labeled astrocytes or adjacent microglia were manually traced, and data were averaged per plaque or image field as a single biological replicate. Parenchymal Aβ burden was quantified as the total Aβ-positive area per section, while vascular Aβ deposition was measured as the Aβ-positive area normalized to CD31⁺ vascular length (μm), calculated using ImageJ. More than 20 sections per group from at least four mice were analyzed for each condition. Quantitative values were normalized to APOE3-expressing controls (set as 1.0) for comparative analysis across groups.

### Transcriptomic profiling

Cortical and hippocampal tissues from AAV-injected mice were microdissected and processed for total RNA extraction using a kit (Vazyme, RC112-01). RNA integrity was verified using an Agilent 2100 Bioanalyzer. Library preparation and paired-end RNA sequencing were performed by Novogene (Beijing, China). Raw sequencing reads (FASTQ files) were quality-checked using FastQC (v0.12.1) and trimmed with Trim Galore (v0.6.10) to remove adapter sequences and low-quality bases. Cleaned reads were aligned to the GRCm38 mouse reference genome using HISAT2 (v2.2.1), and alignment files (SAM format) were converted and sorted using Samtools (v1.19.2). Gene-level read counts were generated using featureCounts (v2.0.6). Genes with zero counts across all replicates were excluded, and expression values were normalized to transcripts per million (TPM). Differentially expressed genes (DEGs) were identified using two-tailed Student’s *t*-tests in R. Venn diagrams were generated using the VennDiagram package to visualize overlaps across treatment groups. Gene Ontology (GO) enrichment analysis (biological process, molecular function, and cellular component) was performed using clusterProfiler (v4.12.6), and shared GO terms were visualized as dot plots. Genes from overlapping GO categories were intersected with a core list of 75 concordantly regulated genes to generate a Z-score-scaled heatmap using the ComplexHeatmap package in R.

### Quantitative real-time PCR (qPCR)

Brain tissues were homogenized and processed for total RNA extraction using RNA-easy Isolation Reagent (Vazyme), following the manufacturer’s instructions. RNA concentration and purity were determined using a Nanodrop spectrophotometer. To eliminate genomic DNA contamination, 1 μg of total RNA was reverse-transcribed into cDNA using the HiScript® III 1st Strand cDNA Synthesis Kit with gDNA wiper (Vazyme). Quantitative PCR was performed using AceQ® qPCR SYBR Green Master Mix (Low ROX Premixed; Vazyme) on an M6000 Real-Time PCR System (LongGene Scientific Instruments) with gene-specific primers (listed in Supplementary Table 2). Reactions were run in technical triplicates. Relative gene expression was calculated using the 2^–ΔΔCt method, with GAPDH used as the internal reference gene.

### Primary cell cultures

Primary cortical neurons were prepared from embryonic day 16–17 (E16–E17) C57BL/6J mouse embryos. Cerebral cortices were dissected, meninges removed, and tissues dissociated in 0.25% trypsin-EDTA (Gibco) at 37°C for 15 min. After trituration and centrifugation, cells were resuspended in Neurobasal medium (Gibco) supplemented with 2% B27, 0.5 mM GlutaMAX, and 1% penicillin-streptomycin. Cells were plated at a density of 1 × 10^5^ cells/cm^2^ on poly-D-lysine-coated plates or coverslips and maintained at 37°C in a 5% CO₂ incubator. Half-medium changes were performed every 3 days, and experiments were conducted on days in vitro (DIV) 6-7. Primary astrocytes were isolated from the cerebral cortices of postnatal day 1–3 (P1–P3) C57BL/6J mice. Cortices were dissected, minced, and digested with 0.25% trypsin-EDTA for 15 min at 37°C, followed by mechanical trituration. Cells were cultured in DMEM/F12 medium supplemented with 10% fetal bovine serum (FBS; Gibco) and 1% penicillin-streptomycin in T75 flasks coated with poly-D-lysine. Medium was replaced every 2–3 days. After 7–10 days, flasks were shaken at 260 rpm for 2 h at 37°C to remove microglia and oligodendrocyte precursor cells. Adherent astrocytes were trypsinized and replated for experiments. Primary microglia were obtained from the same mixed glial cultures used for astrocytes. After 10–12 days in vitro, flasks were shaken at 260 rpm for 2 h to collect loosely adherent microglia in the supernatant. Cells were pelleted, resuspended in DMEM/F12 with 10% FBS, and plated on poly-D-lysine-coated culture dishes. For RAB24 overexpression, mouse RAB24 cDNA (MiaoLing) was amplified by PCR, fused to a C-terminal Flag tag, and inserted into MCS of the pAAV-GfaABC1D-T2A-MCS vector. Cells cultured in 24-well plates were transduced with AAV-RAB24-Flag at a dose of 1.8 × 10^6^ vg per well. For RAB24 knockdown, a small interfering RNA (siRNA) targeting mouse RAB24 (5′-UGUGGCACCAAGAGUGACCUU-3′) was transfected into cells using Lipo8000 reagent (Beyotime) following the manufacturer’s instructions.

### Aβ localization and clearance assay

Synthetic FAM-labeled or unlabeled Aβ42 peptides (AnaSpec) were pretreated with trifluoroacetic acid and 2-propanol as previously described^20^, aliquoted, and stored at −80 °C. Peptides were freshly dissolved in DMSO to 200 μM immediately prior to use. For Aβ localization analysis, cultured cells were incubated with 1 μM FAM-Aβ42 for 24 h, followed by fixation and immunostaining for LAMP1. Co-localization of FAM-Aβ42 with LAMP1⁺ lysosomes was quantified using ImageJ by measuring overlapping fluorescence signals across multiple randomly selected cells from 3 independent biological replicates. For the Aβ clearance assay, cells were first incubated with 1 μM unlabeled Aβ42 for 2 h to allow uptake. After removal of Aβ-containing medium, cells were washed and incubated in fresh medium for an additional 8 h. Cells were then lysed in 5 M guanidine hydrochloride (in 50 mM Tris-HCl, pH 8.0), and intracellular Aβ42 levels were measured using a human Aβ42 ELISA kit (Invitrogen, KHB3441) according to the manufacturer’s protocol. Aβ42 clearance efficiency was calculated as the reduction in cell-associated Aβ42 levels between the 2 h and 10 h timepoints from 3 independent biological replicates.

### Filipin staining

To assess intracellular cholesterol distribution, cells cultured on coverslips were fixed in 4% PFA for 20 min at room temperature (RT) and rinsed with PBS. Fixed cells were incubated with 1.5 mg/mL glycine in PBS for 10 min to quench residual PFA, followed by permeabilization with 0.1% Triton X-100 in PBS for 10 min. Cells were then stained with filipin III (Sigma-Aldrich, F4767) at 50 μg/mL in PBS for 1 h at RT in the dark. Where indicated, immunofluorescence for lysosomal marker LAMP1 was performed prior to filipin staining. Coverslips were mounted using antifade medium and imaged using a fluorescence microscope. Filipin intensity within LAMP1⁺ lysosomal compartments was quantified using ImageJ by measuring co-localized fluorescence signal across multiple randomly selected cells from 3 independent biological replicates.

### Data and statistical analyses

Data are presented as mean ± standard deviation (SD) from at least three independent biological replicates. Statistical analyses were performed using one-way, two-way analysis of variance (ANOVA) followed by Tukey’s HSD post hoc tests, or student’s t-test where indicated. A p-value < 0.05 was considered statistically significant. All analyses were conducted using GraphPad Prism 8.0 software.

## Acknowledgements

We thank the Public Platform of the State Key Laboratory of Natural Medicines at China Pharmaceutical University for access to analytical instrumentation. We also sincerely thank Jie Zhao and Zhenglin Hao (Animal Experimental Center, China Pharmaceutical University) for their technical support with animal experiments. Schematic illustrations were created using BioRender.

## Author contributions

R.Y., Y.L., and J.S. conceived the project and J.S. supervised the study. R.Y., Y.L., X.Z., L.W., and J.W. performed the experiments. R.Y., Y.L. and J.S. analyzed the data and wrote the manuscript.

