## Supplementary information for "AAV-based temporal APOE4-to-APOE2 replacement reveals rebound adaptation and RAB24-mediated Aβ and cholesterol dysregulation"

### Fig. S1

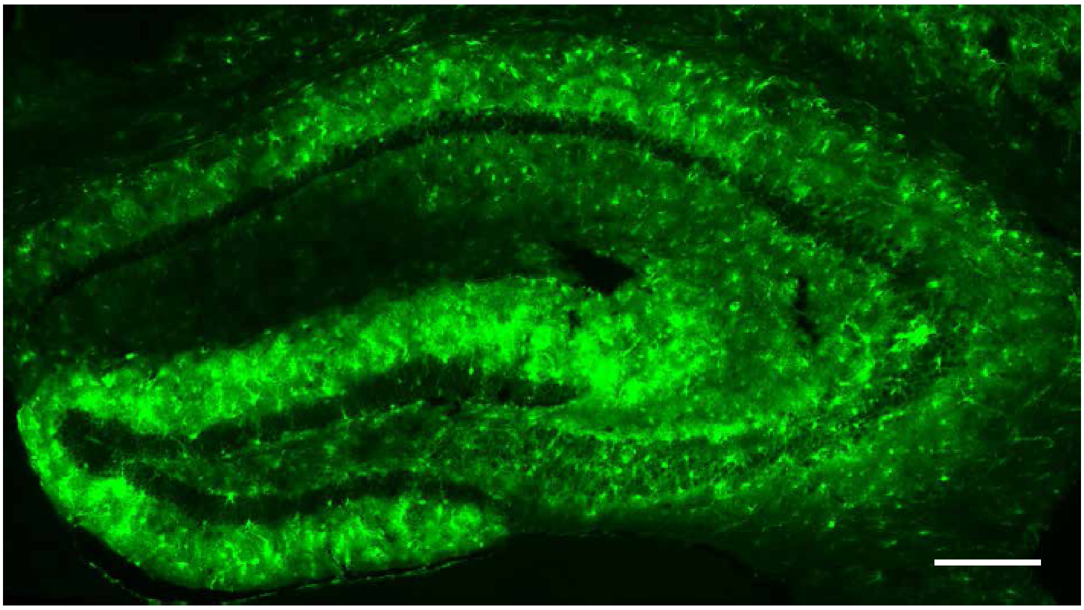

**Fig. S1 Validation of AAV5-mediated transduction efficiency in the wild-type mouse hippocampus**

Representative image showing robust EGFP fluorescence in the hippocampus two weeks after AAV5 injection, confirming efficient viral transduction. Scale bar, 100 μm.

### Fig. S2

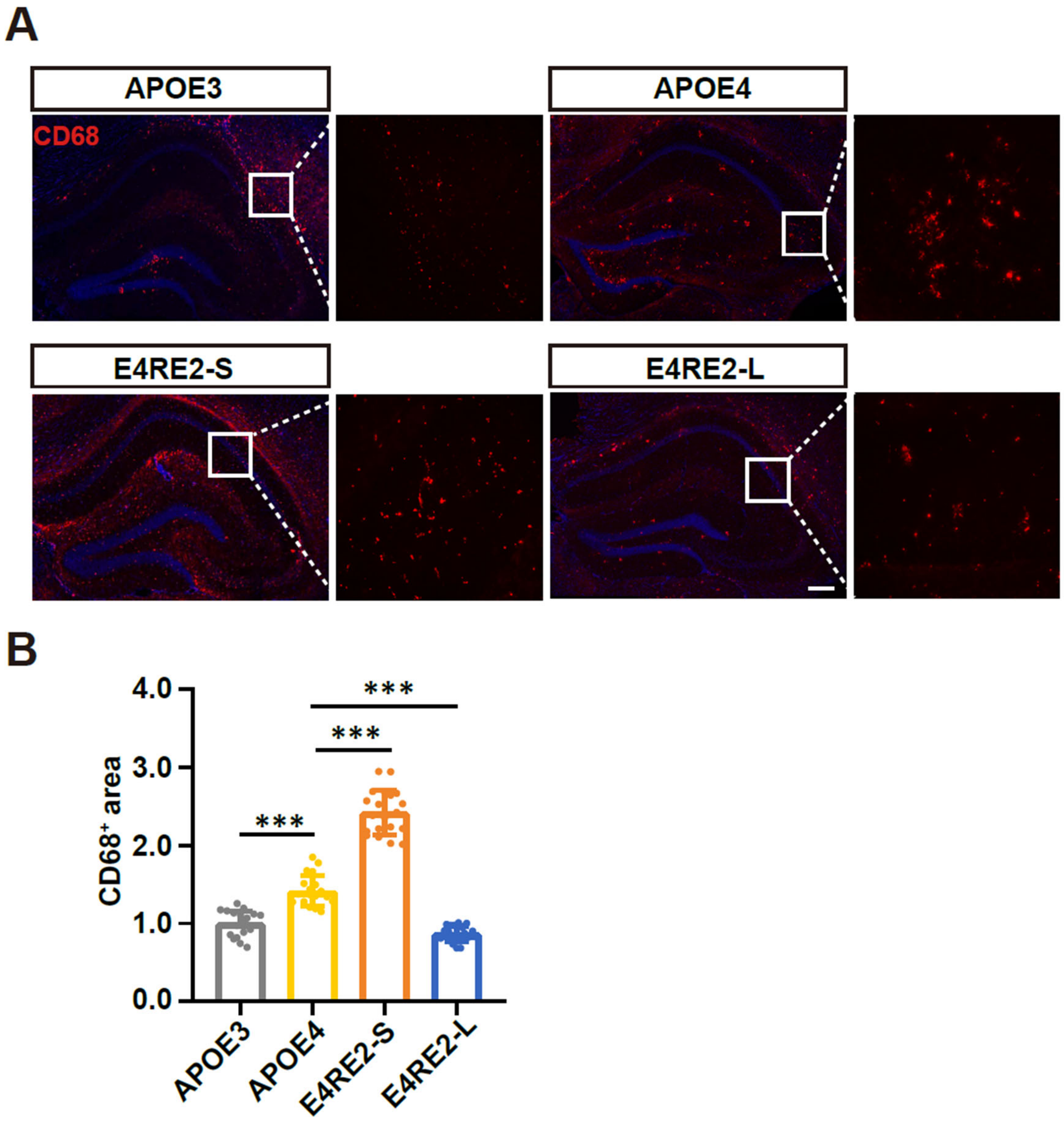


**Fig. S2 Microglial activation is exacerbated by short-term APOE2 replacement in 5×FAD mice**

(A) Representative immunofluorescence images of $CD68^+$ microglia in the hippocampus across four experimental groups: APOE3, APOE4, short-term APOE4-to-APOE2 replacement (E4RE2-S), and long-term replacement (E4RE2-L). (B) Quantification of $CD68^+$ area across treatment groups. Statistical analysis was performed using one-way ANOVA followed by Tukey's HSD post hoc test. Data represent mean ± SD. *$p < 0.05$, **$p < 0.01$, ***$p < 0.001$. Scale bar, 100 μm.

### Fig. S3

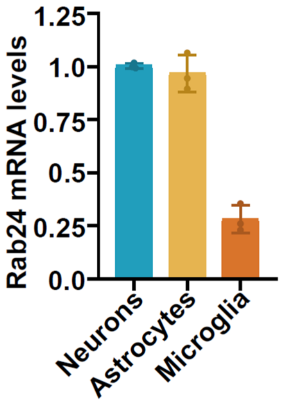


**Fig. S3 RAB24 mRNA expression levels in primary brain cells.**

qPCR analysis of RAB24 mRNA expression levels in primary neurons, astrocytes, and microglia. Expression in neurons was normalized to 1.0. Data are presented as mean ± SD.

**Supplementary Table 1. Antibodies used in this study**

| Antibody | Company | Cat.no. | Dilution |
|---|---|---|---|
| HA-tag | Cell Signaling Technology | 3724 | IF (1:500) |
| Myc-tag | Cell Signaling Technology | 2276S | IF (1:500) |
| GFAP | Cell Signaling Technology | 15913 | IF (1:500) |
| IBA1 | Cell Signaling Technology | 17198 | IF (1:500) |
| CD31 | BD Biosciences | 550274 | IF (1:50) |
| CD68 | Cell Signaling Technology | 97778S | IF (1:500) |
| RAB24 | ProteinTech | 11445-1-AP | IF (1:500) |
| Flag-tag | Sigma | F1804 | IF (1:500) |
| Aβ | Cell Signaling Technology | 2454S | IF (1:500) |
| LAMP1 | DSHB | 1D4B-S | IF (1:100) |
| Goat anti-Rabbit IgG (H+L) Alexa Fluor(R)-568 | Thermo | A11011 | IF (1:500) |
| Goat anti-Rabbit IgG (H+L) Alexa Fluor 633 | Thermo | A21070 | IF (1:500) |
| Goat anti-Rat IgG (H+L) Alexa Fluor™ 546 | Thermo | A11081 | IF (1:500) |
| Goat anti-Rabbit IgG (H+L) Alexa Fluor® 488 | Jackson ImmunoResearch | 167908 | IF (1:500) |
| Goat anti-Mouse IgG (H+L) Alexa Fluor™ Plus 488 | Thermo | A32723 | IF (1:500) |

**Supplementary Table 2. qPCR primers used in this study**

| Primer | sequence |
|---|---|
| Tgfbr1-F | TGCTCCAAACCACAGAGTAGGC |
| Tgfbr1-R | CCCAGAACACTAAGCCCATTGC |
| Tmprss6-F | CTGGTGAGTTCCTCTGCTCTGT |
| Tmprss6-R | TGCTGTCCTCTTGGCACTGGAA |
| Lair1-F | CGCCACTGACGACTCACTC |
| Lair1-R | CTGCCACAAACCAGCAGTTG |
| Rab24-F | GTGGACGTTAAGGTGGTTATGC |
| Rab24-F | CCCGATGGTGTTCTGATAGGG |
| Bcat2-F | ACAGACCACATGCTGATGGTG |
| Bcat2-R | CTGGGTGTAGCGTGAGGTTC |
| Cd72-F | ATGGCTGACGCTATCACGTAT |
| Cd72-R | CCTGTCCTAGATGGTTAGATGCG |
| Mir7228-F | TGGCGACCTGAACAGATG |
| Mir7228-R | GAACATGTCTGCGTATCTC |
| Ring1-F | AGGCCAAAAAGACAAGCGAAA |
| Ring1-R | TTCGGCGCTGTTCCTAAAGG |
| Psmd1-F | CGTTTGGCACCGTTTGGAAG |
| Psmd1-R | TCATCTGTCTAACCACATGGAGA |
| Adcy7-F | GTGCTGGTGTATGTCGAGTG |
| Adcy7-R | GCCTAGTACCATGAGGCAAGC |
| Cav1-F | GCGACCCCAAGCATCTCAA |
| Cav1-R | ATGCCGTCGAAACTGTGTGT |
| Or1e19-F | TCCCCCTTCATTACATGAGCA |
| Or1e19-R | AAGCCACTTTAAGCAGAGTAGAC |
| GAPDH-F | CATCACTGCCACCCAGAAGACTG |
| GAPDH-R | ATGCCAGTGAGCTTCCCGTTCAG |
